# Double-dose mRNA vaccination to SARS-CoV-2 progressively increases recognition of variants-of-concern by Spike RBD-specific memory B cells

**DOI:** 10.1101/2022.08.03.502703

**Authors:** Gemma E. Hartley, Emily S.J. Edwards, Nirupama Varese, Irene Boo, Pei M. Aui, Scott J. Bornheimer, P. Mark Hogarth, Heidi E. Drummer, Robyn E. O’Hehir, Menno C. van Zelm

## Abstract

**Background:** SARS-CoV-2 vaccination with BNT162b2 (Pfizer BioNTech) has been shown to be 95% effective.^1^ Double-dose vaccination generates high levels of spike-specific antibodies, memory B cells (Bmem) and T cells. However, variants of concern (VoC) with mutations in the spike Receptor Binding Domain (RBD) can evade antibody responses. Booster vaccinations improve antibody recognition of VoC, but it is unclear if this is due to higher total antibodies or their capacity to bind VoC. We here addressed the capacity of surface Ig on single Wuhan-specific Bmem after first and second dose BNT162b2 vaccination to recognize variant RBD.

**Methods:** Samples were collected from 30 healthy COVID-19 naive individuals pre-BNT162b2 vaccination, 3 weeks post-dose 1 and 4-weeks post-dose 2. Plasma antibodies and Bmem were evaluated using recombinant RBD proteins of the Wuhan, Gamma and Delta strains.

**Results:** All individuals generated a robust antibody response to BNT162b2 vaccination with all participants producing neutralizing antibodies following dose 2. IgM^+^ and IgG^+^ RBD-specific Bmem were generated after one vaccine dose, and those expressing IgG1 increased in absolute number after dose 2. The majority of RBD-specific Bmem bound the Gamma and/or Delta variants, and this proportion significantly increased after the second dose.

**Conclusion:** The second dose of BNT162b2 increases the number of circulating Ig-class switched RBD-specific Bmem. Importantly, the second dose of vaccination is required for a high frequency of RBD-specific Bmem to recognize Gamma and Delta variants. This suggests that dose 2 not only increases the number of RBD-specific Bmem but also the affinity of the Bmem to overcome the point mutations in VoC.

## INTRODUCTION

The coronavirus disease-2019 (COVID-19) pandemic has now entered its third year. The virus responsible, severe acute respiratory syndrome coronavirus-2 (SARS-CoV-2) has caused ~530 million infections and over 6 million deaths worldwide.^2^ The combined effort of the scientific community has allowed the rapid production and administration of effective SARS-CoV-2 vaccines.^1,3,4^ In developed countries, vaccination rates are high (60-90%), resulting in a reduction of severe disease and hospitalization.^5,6^ In Australia, the ChAdOx1 nCoV-19 (AstraZeneca, adenoviral vector) and BNT162b2 (Pfizer-BioNTech, mRNA) vaccines were widely used for the primary double-dose schedule.^7^ Both the BNT162b2 and mRNA-1273 (Moderna, mRNA) vaccines are now recommended as a 3^rd^ dose booster for all >16 years of age, and a 4^th^ dose for risk groups.^8^

All three vaccines target the SARS-CoV-2 spike protein.^1,3,4^ Antibodies directed towards the Spike receptor binding domain (RBD) prevent binding to the host receptor Angiotensin-converting enzyme 2 (ACE2) and hence can neutralize the virus.^9,10^ After double dose SARS-CoV-2 vaccination, spike-specific antibodies are generated in large quantities peaking between 15-20 days and then begin to decline thereafter as part of the contraction of the immune response.^11–13^

In addition to serum antibodies, SARS-CoV-2 vaccination elicits the formation of Spike-specific memory B cells (Bmem), which predominantly carry surface Ig of either the IgM or IgG isotype.^14,15^ The second vaccine dose is reported to reduce the proportion of IgM^+^ Bmem with a corresponding increase in IgG^+^ Bmem frequencies,^12,14,15^ suggestive of re-activation of pre-existing Bmem in a secondary germinal center (GC) with further affinity maturation following somatic hypermutation (SHM). In humans, CD27^+^IgM^+^IgD^-^ and CD27^-^IgG^+^ Bmem originate predominantly from primary GC reactions, whereas CD27^+^IgG^+^, CD27^+^IgA^+^ and CD27^+^IgE^+^ Bmem display molecular signs of secondary GC responses with higher SHM and increased replication histories.^16–18^

Despite vaccine roll-out in 2021, multiple new variants of concern (VoC) have emerged. VoC are SARS-CoV-2 strains that are deemed by the World Health Organization (WHO) as variants that either increase transmission, disease severity or decrease the effect of public health measures such as vaccination.^19^ VoC Beta (P.1) and Gamma (B.1.351) are mutated at 3 residues within the receptor binding domain (RBD): K417N (Beta) / K417T (Gamma), E484K and N501Y.^19^ The shared E484K mutation results in a 2-6 fold reduction in binding of Wuhan-specific antibodies.^20,21^ In contrast, Delta (B. 1.617) carries two mutations in the RBD: L452R and T478K. Both the E484K and L45R mutations are located within the RBD-2 epitopic region, and can impact antibody neutralization capacity.^22^ Still, antibody recognition of Beta and Gamma are more greatly impacted than Delta.^20,23–27^

In previously-infected individuals, one vaccine dose elicits neutralizing levels of antibody to VoC to the same degree as two doses in naive individuals.^14^ This suggests that re-activation of pre-existing Bmem to undergo a secondary GC response improves recognition of VoCs.^14,27^ However, this has not been addressed in single Bmem cells.

Whilst Australia did not escape community transmission in 2020 and 2021, cases were minimal, and the vast majority of the population was infection-naive during the first year of the vaccine rollout.^28^ Therefore, this population is well-suited to examine Bmem formation after prime and boost vaccination in the absence of SARS-CoV-2 infection. Samples were taken 3-4 weeks following vaccine doses to evaluate the resting Bmem compartment.^29,30^ Using fluorescently-labelled recombinant RBD tetramers, we here examined the SARS-CoV-2-specific Bmem compartment in 30 healthy adults after first and second dose BNT162b2 for their immunophenotype and capacity to bind to the Gamma and Delta variant RBD.

## METHODS

### Participants

Healthy individuals without hematological or immunological disease were enrolled in a low-risk research study to examine their peripheral blood B-cell subsets (Alfred Health ethics no. 32-21/Monash University project no. 72794). From February to June 2021, 30 individuals, who had decided to take the COVID-19 vaccine, consented to three donations of 40ml of blood as well as the collection of basic demographics (age and sex). All individuals were vaccinated with two doses of BNT162b2 (Pfizer-BioNTech) as per the manufacturer’s recommendation. Samples were taken pre-vaccination, 3-4-weeks post-dose 1 and 4-weeks post-dose 2 (range: 19-31 and 25-28 days, respectively) (**Figure 1A**). This study was conducted according to the principles of the Declaration of Helsinki and approved by local human research ethics committees.

**Figure 1.**
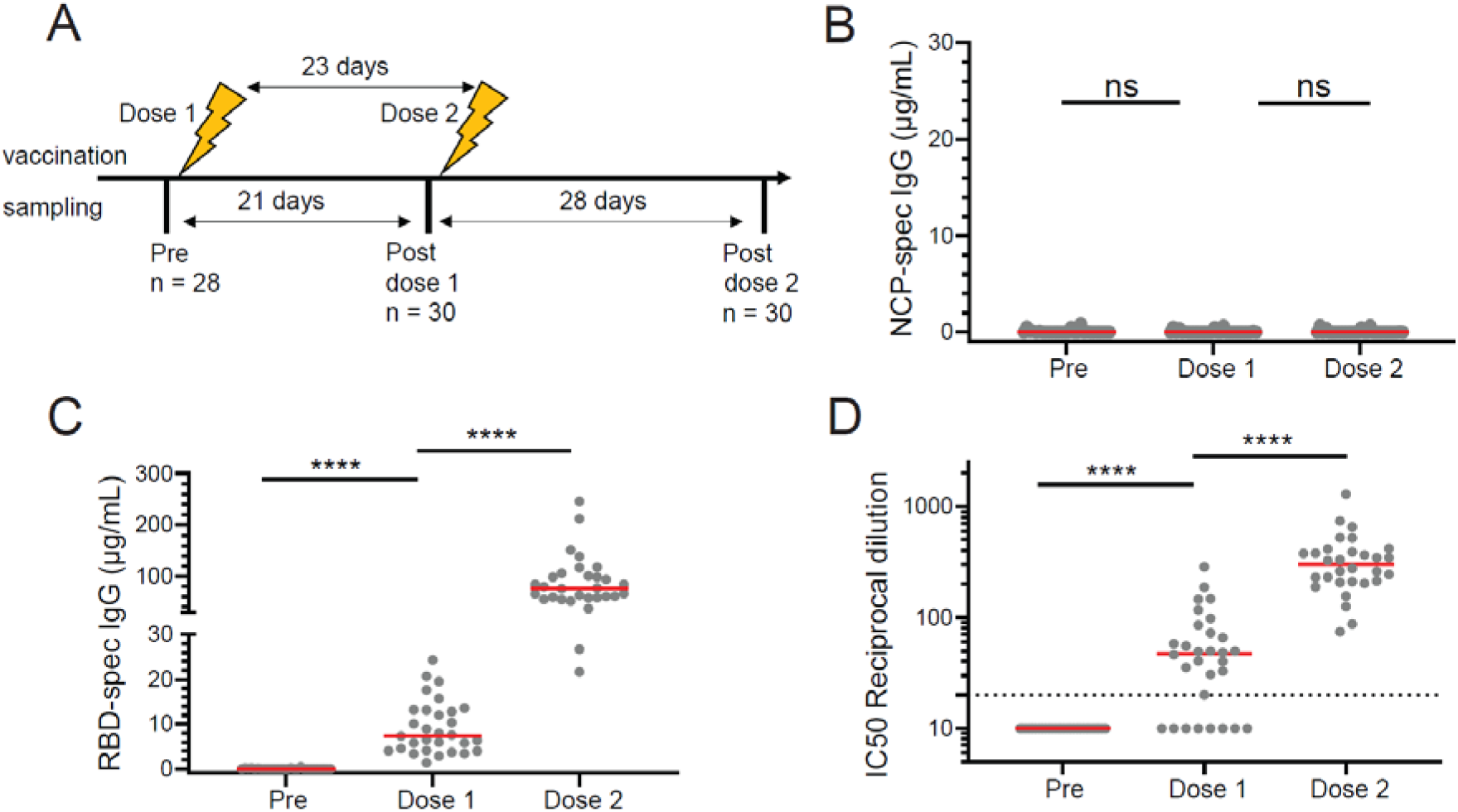
Serological responses BNT162b2 Pfizer vaccination. (A) Participants were sampled pre-BNT162b2 vaccination, 3 weeks after dose 1 and 4 weeks after dose 2. (B) NCP-specific and (C) RBD-specific plasma IgG post-vaccination. (D) Neutralizing antibodies to Wuhan post-vaccination. Dotted line in D depicts IC50 = 20, the cut off for neutralization.^59^ Horizontal lines represent median values. Wilcoxon matched-pairs signed rank test, **** p < 0.0001.

### Sample processing

Blood samples were processed as previously described.^31^ Briefly, 200 μl was used for whole blood cell counts (Cell Dyn analyzer; Abbott Core Laboratory, Abbott Park, IL) and Trucount analysis (see flow cytometry section). The remainder of the sample was used to separate and store plasma (−80°C), and to isolate live peripheral blood mononuclear cells (PBMC) by Ficoll-paque density gradient centrifugation and cryopreservation at a cell density of 10 million cells/ml in RPMI medium with 40% FCS and 10% DMSO in liquid nitrogen for later analysis of RBD-specific B cells.

### Protein production and tetramerization

Recombinant spike RBD proteins of the SARS-CoV-2 Wuhan-1 strain and the Beta, Gamma and Delta variants were produced with the Fel d 1 leader sequence and a biotin ligase (BirA) AviTag target sequence and a 6His affinity tag at the C-terminus, as described previously.^31^ The RBD from the VoC contained the following mutations: B.1.351 (Beta) K417N, E484K, N501Y; P.1 (Gamma) K417K, E484K, N501Y; B.1.617.2 (Delta) L452R, T478K. The DNA constructs were cloned into a pCR3 plasmid and produced and purified, as described previously.^31^ Briefly, plasmid DNA was purified from *E. coli* by Maxiprep (Zymo Research, Irvine, CA), and 30 μg DNA was transfected into 293F cells using the Expi293 Expression system (Thermo Fisher Scientific, Waltham, MA). Supernatants were collected and purified using a Talon NTA-cobalt affinity column (Takara Bio, Kusatsu, Shiga, Japan) with elution in 200 mM Imidazole. Purified proteins were then dialyzed into 10mM Tris and biotinylated, as described previously. Biotinylated protein was subsequently dialyzed against 10 mM Tris for 36 hours at 4° C with minimum of three exchanges, and subsequently stored at −80° C prior to use. Soluble biotinylated RBD Wuhan protein was tetramerized by the addition of either Brilliant Ultra Violet (BUV)395-conjugated streptavidin, or streptavidin-BUV737, and biotinylated RBD Gamma and Delta with streptavidin-BV480 or streptavidin-BV650 (all from BD Biosciences, San Jose, CA) at a protein: streptavidin molar ratio of 4:1 making 4 unique tetramers, [RBD Wuhan]_4_-BUV395, [RBD Wuhan] _4_-BUV737, [RBD Gamma]_4_-BV480 and [RBD Delta]4-BV650.^31,32^

### Measurement of SARS-CoV-2 neutralizing antibodies in plasma

Measurement of neutralizing antibodies was performed using SARS-CoV-2 retroviral pseudotyped particles and a 293T-ACE2 cell line^33^, as described previously. ^31,34^ Briefly, plasma was heat inactivated at 56°C for 45 minutes followed by serial dilution in DMF10. Duplicate serial dilutions were mixed with an equal volume of SARS-CoV-2 (Wuhan-1, Beta, Gamma or Delta spike) retroviral pseudotyped virus and incubated for 1 hour at 37°C. Virus-plasma mixtures were added to 293T-ACE2 cell monolayers seeded the day prior at 10,000 cells/well, and incubated for 2 hours at 37°C before addition of an equal volume of DMF10 and incubated for 3 days. After incubation, tissue culture fluid was removed, monolayers were washed once with PBS and lysed with cell culture lysis reagent (Promega, Madison, WI) and luciferase measured using luciferase substrate (Promega) in a Clariostar plate reader (BMG LabTechnologies, Offenburg, Germany). The percentage entry was calculated as described previously^31^ and plotted against reciprocal plasma dilution GraphPad Prism 9 Software (GraphPad Software, La Jolla, CA) and curves fitted with a one-site specific binding Hill plot. The reciprocal dilution of plasma required to prevent 50% virus entry was calculated from the non-linear regression line (ID50). The lowest amount of neutralizing antibody detectable is a titer of 20. All samples that did not reach 50% neutralization were assigned an arbitrary value of 10.

### ELISA

EIA/RIA plates (Costar, St Louis, MO) were coated with 2 μg/ml recombinant SARS-CoV-2 NCP or RBD overnight at 4° C. Wells were blocked with 3% BSA in PBS and subsequently incubated with plasma samples. Plasma was diluted 1:30 for quantification of RBD- and NCP-specific antibodies pre-vaccination, post-dose 1 and post-dose 2. Plasma was titrated from 1:30 to 1:10,000 for quantification of RBD- and RBD variant-specific antibodies post-dose 1 and 2. Antigen-specific IgG was detected using rabbit anti-human IgG HRP (Dako, Glostrup, Denmark). ELISA plates were developed using TMB solution (Life Technologies, Carlsbad, CA) and the reaction was stopped with 1 M HCl. Absorbance (OD450nm) was measured using a Multiskan Microplate Spectrophotometer (Thermo Fisher Scientific). Serially diluted recombinant human IgG (in-house made human Rituximab) was used for quantification of specific IgG in separate wells on the same plate. Area under the curve (AUC) was calculated for each titration curve using GraphPad Prism software. Relative recognition of the RBD variants was calculated as a percentage of the AUC for that variant relative to the AUC for RBD Wuhan.

### Flow cytometry

Absolute numbers of leukocyte subsets were determined, as previously described. ^31,32,35^ Briefly, 50 μl of whole blood was added to a Trucount tube (BD Biosciences) together with 20 μl of antibody cocktail containing antibodies to CD3, CD4, CD8, CD16, CD19, CD56 and CD45 from the 6-color TBNK reagent kit (BD Biosciences) (**Supplementary Tables 1 and 2**), and incubated for 15 minutes at room temperature in the dark. Subsequently, samples were incubated for a further 15 minutes at room temperature with 500 μl of 1X BD Lysis solution (BD Biosciences) to lyse red blood cells. The tube was then stored in the dark at 4° C for up to 2 hours prior to acquisition on the LSRII or FACSLyric analyzers (BD Biosciences).

For detection of antigen-specific Bmem, 12.5 million PBMC were incubated with fixable viability stain 700 (BD Biosciences), antibodies against CD3, CD19, CD21, CD27, CD38, CD71, IgA, IgD, IgG1, IgG2, IgG3, IgG4, (**Supplementary Tables 1 and 2**) and 5 μg/ml each of [RBD Wuhan]4-BUV395, [RBD Wuhan]4-BUV737, [RBD Gamma]4-BV480 and [RBD Delta]_4_-BV650 for 15 minutes at room temperature in a total volume of 250 μl FACS buffer (0.1% sodium azide, 0.2% BSA in PBS). In addition, 5 million PBMC were similarly incubated with fixable viability stain 700 (BD Biosciences), antibodies against CD3, CD19, CD27 and IgD, plus BUV395-, BUV737-, BV480- and BV650-conjugated streptavidin controls (**Supplementary Tables 1 and 2**). Following staining, cells were washed with FACS buffer, fixed with 2% Paraformaldehyde for 20 minutes at room temperature and washed once more. Following filtration through a 70 μM filter, cells were acquired on the 5-laser BD LSRFortessa X-20 (BD Biosciences). Flow cytometer set-up and calibration was performed using standardized EuroFlow SOPs, as previously described (**Supplementary Tables 3 and 4**).^31,32,35,36^

### Data analysis and statistics

All flow cytometry data were analyzed with FlowJo v10 software (BD Biosciences). Statistical analysis was performed with GraphPad Prism 9 Software (GraphPad Software). Matched pairs were analyzed with the non-parametric Wilcoxon matched pairs signed rank test. Correlations were performed using the non-parametric Spearman’s rank correlation. For all tests, *p* < 0.05 was considered significant.

## RESULTS

### Robust antibody responses in all donors after two doses of BNT162b2

Blood samples were collected from 30 healthy COVID-19 naive individuals before and after first and second dose BNT162b2, which were provided with a median of 23 days between doses (range: 21-39 days). A total of 28 samples were collected pre-vaccination (2 samples missed) and 30 samples were obtained 3 weeks post-dose 1 as well as 4-weeks post-dose 2 (**Figure 1A**). IgG to SARS-CoV-2 nucleocapsid (NCP) and Spike RBD was evaluated in the donors before vaccination with BNT162b2 and after dose 1 and dose 2. As the BNT162b2 vaccine only contains mRNA encoding the spike protein, the presence of NCP-specific IgG would be indicative of previous infection. All participants were SARS-CoV-2 naive: they did not have NCP-specific antibodies at commencement of the study and remained negative throughout (**Figure 1B**). None of the participants had detectable IgG to Spike RBD pre-vaccination, further confirming that all donors were SARS-CoV-2 naive (**Figure 1C**). All participants generated anti-RBD IgG post-dose 1, and these levels were significantly increased post-dose 2 (**Figure 1C**). Similarly, none of the participants had neutralizing antibody levels before vaccination, as evaluated using a pseudotyped viral assay (**Figure 1D**).^31^ Most individuals (22/30) generated neutralizing antibodies after the first vaccine dose, and after the second dose all participants carried levels of neutralizing antibodies above an IC50 of 100 (**Figure 1D**). Thus, BNT162b2 vaccination generates high levels of RBD-specific IgG and neutralizing antibodies 4 weeks after dose 2.

### Expansion of RBD-specific Bmem expressing IgG1 after two vaccine doses

To examine RBD-specific Bmem generated towards BNT162b2 vaccination, recombinant RBD-Wuhan protein was biotinylated and tetramerized with fluorescently-labeled streptavidins. Two tetramers were used to identify RBD-specific B cells: [RBD Wuhan]_4_-BUV395 and [RBD Wuhan]_4_-BUV737.^31^ These tetramers were used together with a cocktail of surface antibodies to identify and extensively immunophenotype total and RBD-specific Bmem (**Figure 2A, Supplementary Figure 1A**). Within total CD19^+^ B cells and RBD-specific B cells, naive B cells were defined as IgD^+^CD27^-^ and excluded from further analysis, with the remainder of the population deemed Bmem (**Figure 2A, Supplementary Figure 1A**). IgD^-^ Bmem were then further immunophenotyped using surface antibodies against IgG1,2,3,4 and IgA (**Figure 2A)**. RBD-specific Bmem were detected in all donors after first and second dose vaccination, with significantly higher numbers post-dose 2 (**Figure 2B**). Donor age did not appear to affect the generation of RBD-specific Bmem as older individuals generated similar numbers of cells to younger individuals in the cohort (**Supplementary Figure 2**). The RBD-specific Bmem compartment consisted mostly of IgM^+^ or IgG1^+^ cells (**Figure 2C**). In contrast, the total Bmem population contained fewer IgG1^+^ cells and more IgG2^+^ than the RBD-specific Bmem (**Figure 2C, Supplementary Figure 1B**). After dose 2, RBD-specific IgG1, IgG2 and IgG3 Bmem populations were significantly expanded whereas the total Bmem compartment remained unchanged (**Figure 2C, Supplementary Figures 1B and 2**). The total numbers of RBD-specific IgG^+^ Bmem were positively correlated with RBD-specific plasma IgG after both vaccine doses (**Figure 2D, Supplementary Figure 3**). While the two IgM^+^ Bmem subsets (CD27^+^ IgM^+^ IgD^+^ and CD27^+^IgM^+^ only) were proportionally reduced after dose 2, their absolute numbers were similar to dose 1 (**Figure 2C**). Within the total Bmem compartment, no changes were observed between the samples obtained after dose 1 vs dose 2 (**Supplementary Figure 1B**). Thus, BNT162b2 vaccination specifically affects the antigen-specific Bmem, and the second dose boosts the formation of IgG1^+^ Bmem.

**Figure 2.**
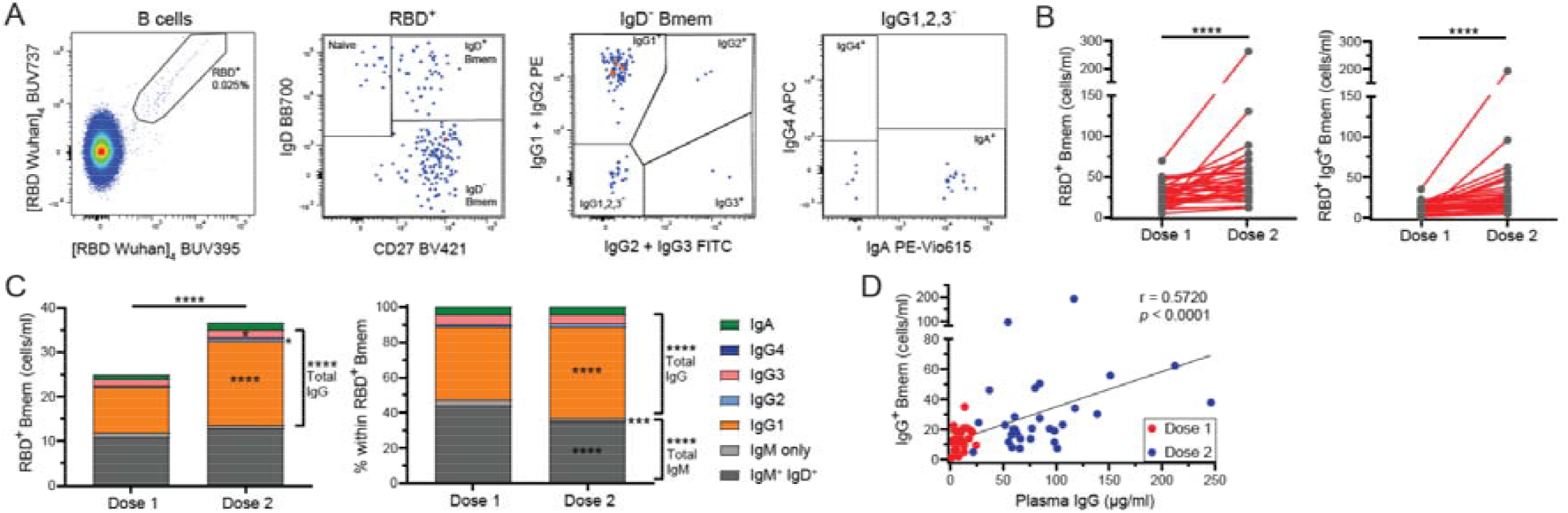
Identification of RBD-specific Bmem elicited by BNT162b2 vaccination. (A) Gating strategy and immunophenotype of RBD-specific memory B cells (Bmem) using RBD Wuhan tetramers. (B) Absolute number of RBD-specific Bmem and IgG^+^ Bmem in vaccinated individuals. (C) Absolute number and proportion of Ig switched RBD-specific Bmem. Wilcoxon matched-pairs signed rank test, * p < 0.05, *** p < 0.001, **** p < 0.0001. (D) Correlation between plasma IgG and IgG+ RBD-specific Bmem after dose 1 and dose 2. Trend line depicts linear correlations; statistics, nonparametric Spearman’s rank correlation.

### RBD-specific Bmem exhibit a resting memory phenotype 3-4 weeks after vaccination

The nature of RBD-specific Bmem was further examined on the basis of CD21, CD27 and CD71 expression (**Figure 3A**). CD27 marks a population of IgG^+^ Bmem with higher antigen-driven replication and signs of antibody maturation than the IgG^-^ counterpart.^16^ Approximately 95% of RBD-specific IgG^+^ Bmem expressed CD27, and this proportion was not different between first and second dose vaccination (**Figure 3C**). Low expression of CD21 marks recently activated Bmem, which were shown to expand after influenza vaccination followed by contraction at week 4.^29^ After BNT162b2 dose 1, approximately 15% of the RBD-specific Bmem were CD21^lo^ This proportion was slightly, but significantly, lower (to 10%) 4 weeks after the second dose (**Figure 3C**). CD71 is another marker of recent Bmem activation, typically expressed after 1-2 weeks and downregulated by week 4.^30^ Approximately 10% of RBD-specific Bmem expressed CD71 both after dose 1 and after dose 2. Within the total Bmem population, no significant differences were observed post-dose 1 vs post-dose 2, demonstrating the stability of the overall Bmem compartment (**Supplementary Figure 1**). Thus, we have established that 4 weeks after BNT162b2 vaccination, RBD-specific Bmem display a resting, mature Bmem immunophenotype.

**Figure 3.**
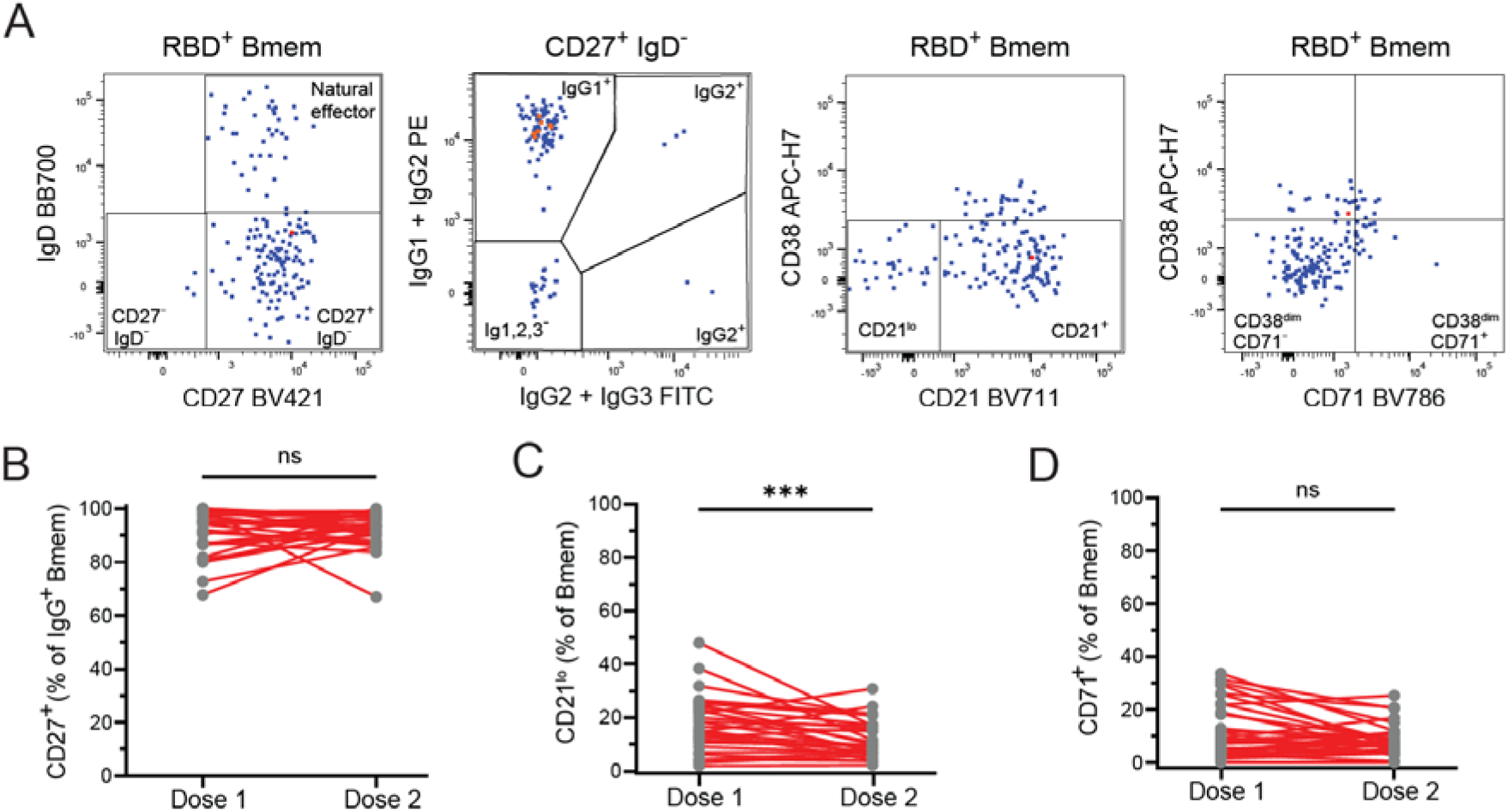
RBD-specific Bmem exhibit a resting phenotype 4 weeks after dose 2 of BNT162b2 vaccination. (A) Gating strategy to delineate CD27^+^, CD21^lo^ and CD71^+^ RBD-specific Bmem. (B) Frequency of RBD-specific IgG^+^ Bmem that express CD27. (C) Frequency of CD21^lo^ RBD-specific Bmem. (D) Frequency of CD71^+^ RBD-specific Bmem. Wilcoxon matched-pairs signed rank test, *** p<0.001.

### The second vaccine dose boosts the capacity of RBD-specific Bmem to recognize SARS-CoV-2 variants of concern

With the first BNT162b2 dose already inducing plasma antibodies and a resting, mature Bmem population, which are further expanded after dose 2, we examined the capacity of these cells to recognize VoC with antibody evasion mutations: Beta and Gamma (E484K) and Delta (L452R) (**Figure 4A**).^37^ Vaccine-induced antibodies were evaluated for their ability to bind variants through ELISA and the pseudotyped neutralization assay.^31^ RBD Wuhan-specific plasma IgG showed partial recognition against all three variants with a more profound decrease observed for Beta and Gamma than for Delta (**Figure 4B**). The second dose of BNT162b2 significantly increased the proportion of Wuhan RBD-specific antibodies able to bind Beta and Gamma (**Figure 4B**). Still the capacity of vaccine generated antibodies to neutralize Beta and Gamma was lower than for the Delta variant, supporting the ELISA results (**Figure 4C**). To evaluate the recognition of RBD-specific Bmem to VoC, tetramers of RBD Gamma and RBD Delta were generated. Combined staining with fluorochrome-conjugated RBD Wuhan tetramers enabled the RBD-specific cells that bound to either RBD Gamma, Delta or neither variant (**Figure 4D**). The Gamma and Delta RBD were each recognized by approximately 50% of RBD-specific Bmem. After the second dose these proportions both significantly increased to 70% (**Figure 4E**). The increased proportion of RBD-specific Bmem that recognized either or both variants were mainly due to an increase in IgG1^+^ Bmem (**Supplementary Figure 4**).

**Figure 4.**
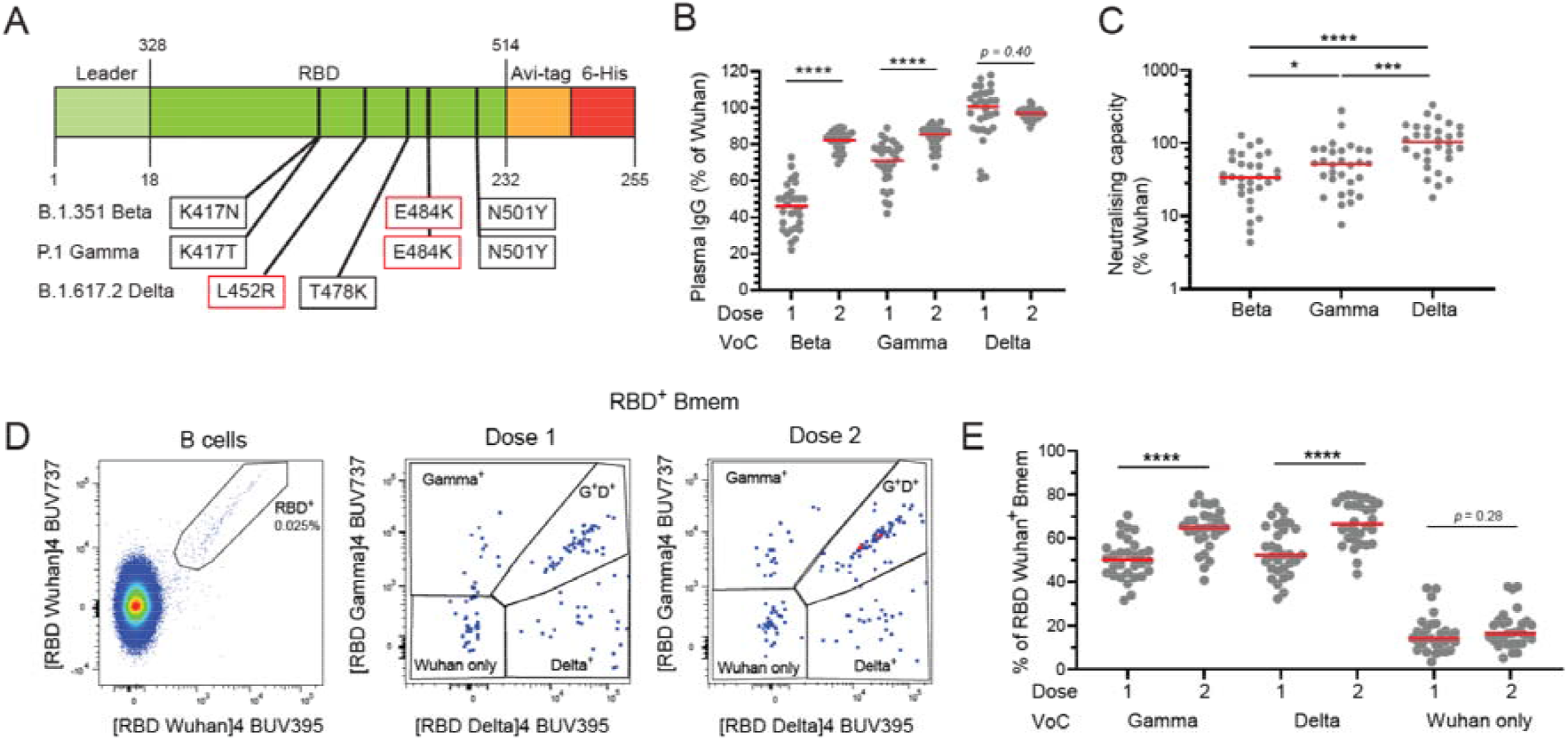
Second BNT162b2 dose increases recognition of variants of concern by RBD-specific antibodies and Bmem. (A) Schematic of point mutations in the RBD for Beta, Gamma and Delta variants. Mutations in red cause a reduction in antibody recognition.^22,37^ (B) Proportion of RBD-specific IgG that also bind VoC Beta, Gamma and Delta after dose 1 and 2. (C) Neutralizing capacity of Wuhan spike-specific antibodies against Beta, Gamma and Delta VoC. (D) Gating strategy to identify RBD-specific Bmem that also bind RBD Gamma and/or Delta. (E) Proportion of RBD-specific Bmem that also bind Gamma, Delta or no variants. Wilcoxon matched-pairs signed rank test, * p < 0.05, *** p < 0.001, **** p < 0.0001.

In summary, double-dose BNT162b2 vaccination effectively elicits RBD-specific IgG, SARS-CoV-2 neutralizing antibodies and RBD-specific Bmem. The second dose enhances the response quantitatively, and improves the capacity of antibodies and Bmem cells to bind mutated RBD domains from the VoC.

## DISCUSSION

We show herein that the first dose of BNT162b2 vaccination robustly induces SARS-CoV-2-specific plasma antibodies and Bmem. RBD-specific Bmem are expanded after dose 2 and display an enhanced capacity to bind to VoC.

The production of RBD-specific and neutralizing antibodies following BNT162b2 vaccination has been well-characterized showing rapid production of antibodies within 1 month of vaccination.^12,14,15,27,38^ Interestingly, not all individuals in our cohort generate neutralizing antibodies after one dose of vaccine, but do so after the second dose. This observation recapitulates results from other cohorts, and demonstrates the need for the double-dose primary schedule.^12,14^ Neutralizing antibodies prevent ACE2 binding and viral entry into host cells,^9,10^ and the levels correlate strongly with vaccine effectiveness.^39^ However, it remains unclear whether serum antibodies are representative of those in the upper airways, the entry site for SARS-CoV-2. Therefore, serum neutralizing antibodies may not be an accurate marker of vaccine protection and effectiveness. Furthermore, as levels of neutralizing antibodies decline beyond 1-month post-vaccination,^11–15^ these do not represent the durable protection from severe disease that lasts 3-6 months.^39^

In contrast to plasma Ig, SARS-CoV-2-specific Bmem are maintained in stable number after infection and vaccination.^15,40^ We here show that the first vaccine dose elicits the formation of RBD-specific Bmem with a resting phenotype (CD27^+^CD71^-^) after 3 weeks, i.e. prior to the administration of the second dose of BNT162b2. The second dose likely induces re-activation of these pre-existing Bmem and differentiation into plasmablasts responsible for the rise in RBD-specific plasma IgG and neutralizing antibodies. This indicates that vaccine-induced RBD-specific Bmem are functional and may contribute to protection from severe disease after infection. Other studies have also shown a predominant IgG^+^ Bmem response after two SARS-CoV-2 vaccine doses, in which the proportion of IgG^+^ Bmem increased with a reciprocal decrease in IgM^+^ Bmem frequencies.^12,14^ We here expand on this by showing that the Bmem response to the second dose of BNT162b2 vaccination is dominated by IgG1^+^ cells. Furthermore, through analysis of absolute cell numbers per milliliter of blood, we show that this is an absolute expansion of IgG1^+^ RBD-specific Bmem, while IgM^+^ Bmem numbers remain at a similar level. Importantly, we note that IgA expressing Bmem are formed in lower frequency and number compared to SARS-CoV-2 infection. ^31^ However, based on the study design of sampling peripheral blood it is difficult to interpret their impact in mucosal upper airway tissues.

The predominance of RBD-specific Bmem expressing IgG1 reflects the serological response, which has been shown to be dominated by plasma IgG1 antibodies. We also show that the number of RBD-specific Bmem positively correlate with RBD-specific IgG in the plasma. The presence of high numbers of IgG1^+^ Bmem also indicates that the second dose re-activates pre-existing Bmem to not only form plasmablasts, but also directs cells to re-enter a GC and undergo further Ig class switching and affinity maturation. GC reactions following SARS-CoV-2 vaccination are maintained up to 7 months post-vaccination, and are thought to drive the observed gradual rise in SHM levels.^41^ It would therefore be of interest for future studies to examine longitudinal samples beyond 6 months post-vaccination, or after booster doses for ongoing class switching to downstream Ig genes (ie. IgG2 and IgG4)^17^ and further affinity maturation.

The question remains, how durable are the numbers of SARS-CoV-2-specific Bmem following adenoviral vaccination? While recent studies have examined the protection against this vaccine formulation,^42–44^ the longevity of this protection is yet to be fully characterized. Antibody responses to two doses of adenoviral vaccination are significantly lower compared to mRNA vaccination.^42,45–47^ However, one dose of ChAdOx1 nCoV-19 followed by one dose of BNT162b2 generates similar antibody levels to two doses of BNT162b2.^43–45,48^ Therefore, an mRNA booster vaccination in individuals that received primary adenoviral vector vaccination provides the same protection as a third dose of mRNA vaccine.^47,49–51^ Durability following mRNA vaccination has been more extensively characterized. Spike-specific Bmem remain in stable frequencies for up to 6 months post-vaccination.^12,14,15^ Responses following a third dose further increases the frequency of these cells and also their ability to bind variants.^40,52^

VoC are now a major threat to the protection that COVID-19 vaccines provide. Mutated RBD proteins in our study, representing the Beta, Gamma and Delta VoC, all demonstrated reduced antibody recognition. ^20,21,23–25^ Hence, we can use the antibody and Bmem cell responses to these VoC as a model of variants that partially escape the current Wuhan based BNT162b2 vaccination. We show that the second vaccine dose increased the capacity of antibodies and Bmem to bind VoC. This suggests that the second dose of vaccination increases the affinity of Bmem, enabling these to overcome the minor changes in the RBD and still bind variant RBDs with sufficient affinity. This is in agreement with other observations that Ig genes from variant-binding Bmem have higher SHM levels than those that only bind to Wuhan^15^ However, with the high number of RBD mutations in the current Omicron (B.1.1.529) subvariants, the vaccine-elicited Bmem might not have the capacity to overcome the mismatches.^40,53–55^ Still, a third-dose booster vaccination with the Wuhan Spike has been shown to improve antibody and Bmem binding to Omicron BA.1.^40^ Thus, the current vaccine formulation does appear to provide protection that improves with multiple doses.^40^ Importantly, Omicron breakthrough responses after double-dose vaccination do not elicit greater capacities of antibodies or Bmem to bind the Omicron Spike protein than Wuhan vaccination.^54,56^ Furthermore, all Bmem after breakthrough infection bind Wuhan with greater affinity than Omicron, indicating that a variant infection does not elicit Bmem with new specificities, ie. Original antigenic sin.^57,58^ However, we have shown that repeat exposure double dose vaccination increases the capacity of Wuhan-specific Bmem to recognize VoC.

It will be critical to examine the protection the BNT162b2 vaccination provides against emerging VoC. Furthermore, if new VoC emerge, current SARS-CoV-2 vaccines may need to incorporate variant epitopes that will elicit new variant-specific antibodies and Bmem. Overall, we have shown that BNT162b2 vaccination generates a strong antibody and Bmem response. The second dose of BNT162b2 vaccine is required for increased recognition of VoC. As VoC are the dominant strains across the globe, it is critical to ensure that multiple dose SARS-CoV-2 vaccinations are continued, and to monitor their ability to protect against severe disease caused by new variants.

## ACKNOWLEDGEMENTS

We thank Dr. Bruce D. Wines and Ms. Sandra Esparon (Burnet Institute) for technical assistance, Mr. Jack Edwards and Ms. Ebony Blight (Monash University) for sample collection and preparation, and the staff of ARAFlowcore for flow cytometry support. Supported by an Australian Government Medical Research Future Fund (MRFF, Project no. 2016108; MCvZ, HED and REO’H) and an unrestricted research grant from BD.

## AUTHOR CONTRIBUTIONS

Designed and/or performed experiments: GEH, ESJE, NV, IB, PMA, PMH, HED, REO’H and MCvZ; Formal analysis: GEH, NV, IB and PMA; Provided reagents: SJB; Supervised the work: ESJE, REO’H and MCvZ; Wrote the manuscript: GEH and MCvZ. All authors edited and approved the final version of the manuscript.

