## Supplemental Material for "Double-dose mRNA vaccination to SARS-CoV-2 progressively increases recognition of variants-of-concern by Spike RBD-specific memory B cells"

**Supplementary Tables (n= 4) and Figures (n= 4)**

**Supplementary Table 1. Composition of the antibody panels**

|  | **Fluorochrome** | | | | | | | | | | | | | | | | |
| --- | --- | --- | --- | --- | --- | --- | --- | --- | --- | --- | --- | --- | --- | --- | --- | --- | --- |
| Tube | **BUV395** | **BUV496** | **BUV737** | **BV421** | **BV480** | **BV605** | **BV650** | **BV711** | **BV786** | **FITC** | **PerCP-Cy5.5/BB700** | **PE** | **PE-Vio615** | **PC7/**  **PE-Cy7** | **APC** | **AF700** | **APC-H7** |
| 1. TruCount | **-** | **-** | **-** | **-** | **-** | **-** |  | **-** | - | CD3 | CD45 | CD16 + CD56 | **-** | CD4 | CD19 | - | CD8A |
| 2. Ag-specific Bmem | RBD Wuhan | CD3 | RBD Wuhan | CD27 | RBD gamma | - | RBD delta | CD21 | CD71 | IgG2 + IgG3 | IgD | IgG1 + IgG2 | IgA | CD19 | IgG4 | Viability | CD38 |
| 3. Streptavidin control | Strep | - | Strep | CD27 | Strep | - | Strep | - | - | CD3 | IgD | - | - | CD19 | - | Viability | - |

**Supplementary Table 2. Antibody list**

| **Marker** | **Fluorochrome** | **Clone** | **Source** | **Cat. number** | | **Volume/**  **test (μl)** | **Tube(s)** |
| --- | --- | --- | --- | --- | --- | --- | --- |
| CD3 | BUV496 | UCHT1 | BD Bioscience | 612940 | 1 | | 2 |
| CD3 | FITC | UCHT1 | BD Biosciences | 555332 | 3 / 1 | | 1 / 3 |
| CD4 | PC7 | SFCI12T4D11 | Beckman Coulter | 6607101 | 0.2 | | 1 |
| CD8A | APC-H7 | SK1 | BD Biosciences | 560179 | 4 | | 1 |
| CD16 | PE | B73.1 | Biolegend | 360704 | 0.2 | | 1 |
| CD19 | APC | SJ25C1 | Biolegend | 363006 | 0.4 | | 1 |
| CD19 | PE-CY7 | SJ25C1 | BD Biosciences | 557835 | 5 | | 2 |
| CD21 | BV711 | B-ly4 | BD Biosciences | 563163 | 5 | | 2 |
| CD27 | BV421 | M-T271 | BD Biosciences | 562513 | 1 | | 2 / 3 |
| CD38 | APC-H7 | HB7 | BD Biosciences | 303534 | 1 | | 2 |
| CD45 | PerCP-Cy5.5 | 2D1 | BD Biosciences | 340953 | 2 | | 1 |
| CD56 | PE | B159 | BD Biosciences | 555516 | 5 | | 1 |
| CD71 | BV786 | M-A712 | BD Biosciences | 563768 | 1 | | 2 |
| IgA | PE-Vio615 | REA1014 | Miltenyi Biotec | 130-116-882 | 1.5 | | 2 |
| IgD | BB700 | IA6-2 | BD Biosciences | 566538 | 1 | | 2 |
| IgG1 | PE | G17-1 | BD Biosciences | 624049 | 0.1 | | 2 |
| IgG2 | PE | HP6002 | BD Biosciences | 624049 | 0.5 | | 2 |
| IgG2 | FITC | HP6002 | BD Biosciences | 624045 | 1 | | 2 |
| IgG3 | FITC | HP6047 | BD Biosciences | 624045 | 0.5 | | 2 |
| IgG4 | APC | SAG4 | Cytognos | CYT-IGG4AP | 2 | | 2 |
| Strep | BUV395 | - | BD Biosciences | 564176 | 0.67 | | 3 |
| Strep | BUV737 | - | BD Biosciences | 564293 | 0.67 | | 3 |
| Strep | BV480 | - | BD Biosciences | 564876 | 0.67 | | 3 |
| Strep | BV650 | - | Biolegend | 405232 | 0.13 | | 3 |
| Viability | AF700 | - | BD Biosciences | 564997 | 0.1 | | 2 / 3 |

**Supplementary Table 3. Flow cytometer set-up**

| **LSRFortessa X-20** | | **LSRII** | | **FACSLyric** | | **Fluorochromes used in this study** |
| --- | --- | --- | --- | --- | --- | --- |
| **355 nm** | | **-** | | **-** | |  |
| 379/28 | No LP | **-** | - | - | - | BUV395 |
| 525/50 | 505 LP | **-** | - | - | - | BUV496 |
| 740/35 | 690 LP | **-** | - | - | - | BUV737 |
| **405 nm** | | **405 nm** | | **405 nm** | |  |
| 450/50 | No LP | 450/50 | No LP | 448/45 | 448/45 | BV421 |
| 525/50 | 505 LP | 525/50 | 505 LP | 528/45 | 500 LP | BV480 |
| - | - | 586/15 | 570 LP | - | - | - |
| 610/20 | 600 LP | 610/20 | 600 LP | 606/36 | 606/36 | - |
| 670/30 | 635 LP | 660/20 | 630 LP | - | - | BV650 |
| 710/50 | 685 LP | 710/50 | 685 LP | 715/50 | 715/50 | BV711 |
| 780/60 | 750 LP | 780/60 | 750 LP | 755 LP | 755 LP | BV786 |
| **488 nm** |  | **488 nm** |  | **488 nm** | |  |
| 488/10 | No LP | 488/10 | No LP | 488/15 | No LP | SSC |
| 530/30 | 505 LP | 530/30 | 505 LP | 527/32 | 507 LP | FITC |
|  |  |  |  | 586/42 | 560 LP | PE |
| 710/50 | 685 LP | 710/50 | 630 LP | 700/54 | 665 LP | PerCP-Cy5.5, BB700 |
|  |  |  |  | 783/56 | 752 LP | PE-Cy7 |
| **561 nm** |  | **561 nm** |  | **-** | |  |
| 586/15 | No LP | 582/15 | No LP | - | - | PE |
| 610/20 | 600 LP | 610/20 | 600 LP | - | - | PE-Vio615 |
| 675/50 | 635 LP | 685/35 | 635 LP | - | - | - |
| 780/60 | 750 LP | 780/60 | 750 LP | - | - | PE-Cy7, PC7 |
| **640 nm** |  | **640 nm** |  | **640 nm** | |  |
| 670/30 | No LP | 670/14 | No LP | 660/10 | 660/10 | APC |
| 730/45 | 690 LP | 730/45 | 690 LP | 720/30 | 705 LP | Fixable Viability Stain 700 |
| 780/60 | 750 LP | 780/60 | 750 LP | 783/56 | 752 LP | APC-H7, APC-Cy7 |

**Supplementary Table 4. Target values for 7^th^ peak of rainbow beads in fluorescent channels**

| **Fluorochrome** | **Channel** | Lower (-15%) | **Target MFI** | Upper (+15%) | Recommendation |
| --- | --- | --- | --- | --- | --- |
| **BUV395** | **UV395** | 17,000 | **20,000** | 23,000 | In-house |
| **BUV496** | **UV525** | 23,800 | **28,000** | 32,200 | In-house |
| **BUV737** | **UV737** | 21,2500 | **25,000** | 28,750 | In-house |
| **BV421** | **V450** | 100,452 | **118,178** | 135,905 | EuroFlow |
| **BV480** | **V525** | 93,871 | **110,436** | 127,002 | EuroFlow |
| **BV650** | **V670** | 34,383 | **40,450** | 46,518 | In-house |
| **BV711** | **V710** | 15,079 | **17,740** | 20,401 | In-house |
| **BV786** | **V780** | 1,903 | **2,239** | 2,575 | In-house |
| **FITC** | **B530** | 28,752 | **33,826** | 38,900 | EuroFlow |
| **PerCP-Cy5.5/ BB700** | **B710** | 66,846 | **78,642** | 90,438 | EuroFlow |
| **PE** | **YG586** | 32,381 | **38,095** | 43,809 | EuroFlow |
| **PE-Vio615** | **YG610** | 178,500 | **210,000** | 241,500 | In-house |
| **PE-Cy7** | **YG780** | 8,316 | **9,783** | 11,250 | EuroFlow |
| **APC** | **R670** | 158,639 | **186,634** | 214,629 | EuroFlow |
| **AF700** | **R730** | 121,550 | **143,000** | 164,450 | In-house |
| **APC-H7** | **R780** | 64,194 | **75,522** | 86,850 | EuroFlow |
| Spherotech Rainbow Calibration particles (8 peaks) 3.41µm; cat nr. RCP-30-5A, Lot No. EAG01  As per EuroFlow recommendation (Kalina *et al.* 2012) | | | | | |

**SUPPLEMENTARY FIGURES (n=4)**

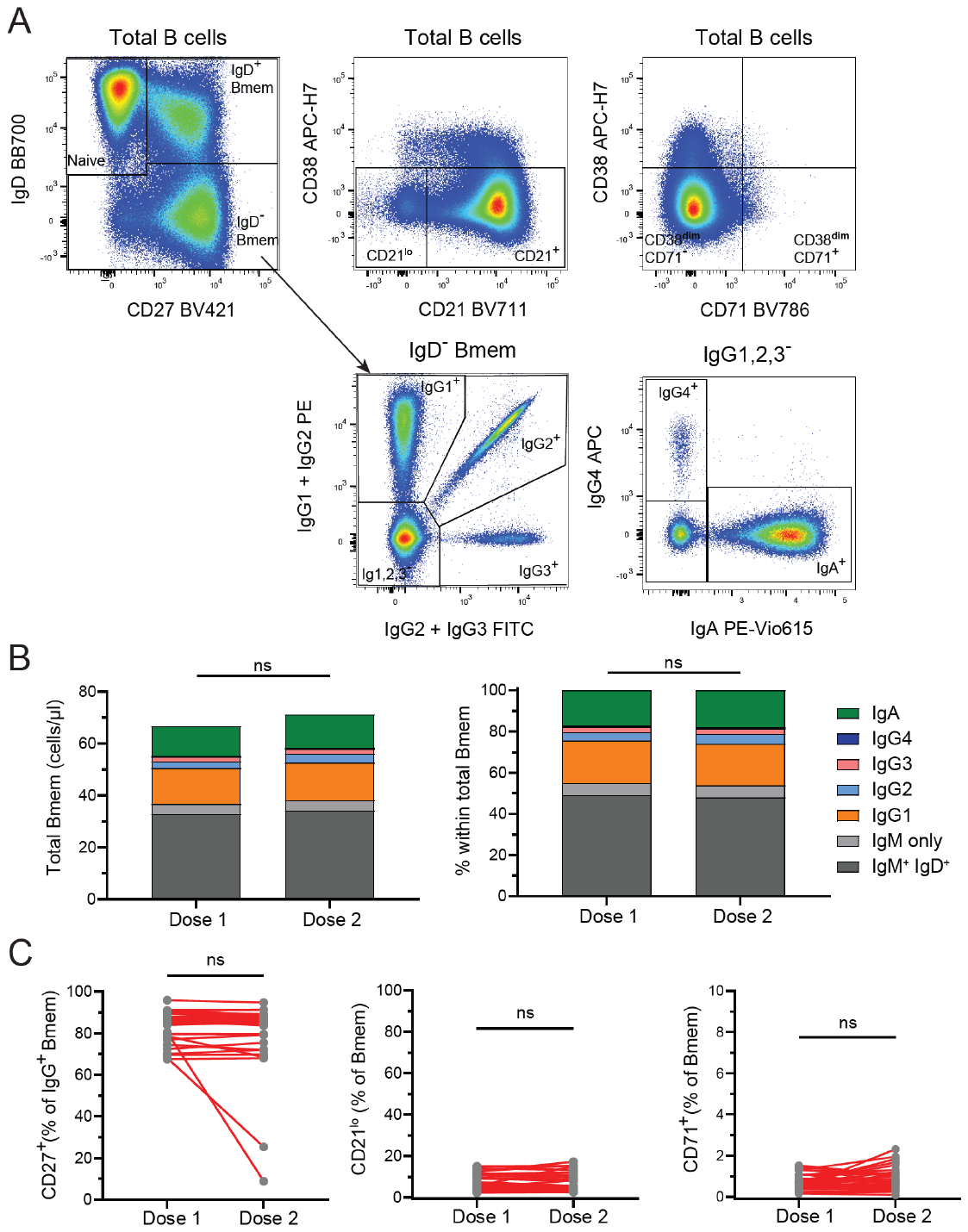

**Supplementary Figure 1. Total Bmem phenotype and Ig distribution post-dose 1 and post-dose 2.** (A) Identification of total Bmem, CD21^lo^, CD71^+^, and Ig switched total Bmem. (B) Absolute number and frequency of total Bmem subpopulations. (C) Frequency of CD27^+^ IgG^+^, CD21^lo^ and CD71^+^ total Bmem. Wilcoxon matched-pairs signed rank test.

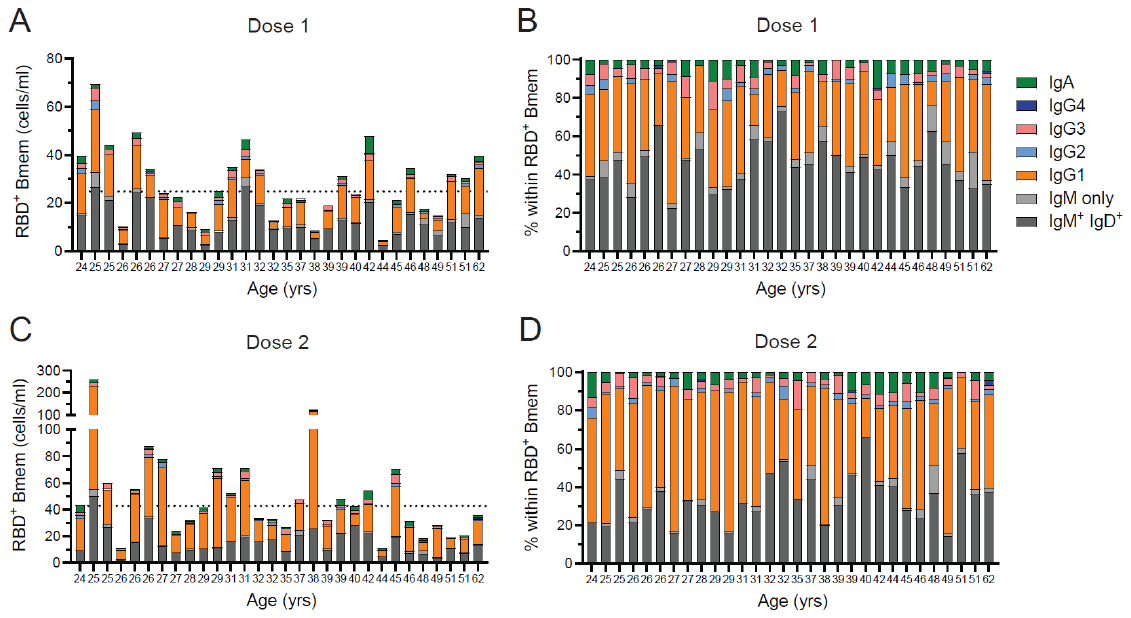

**Supplementary Figure 2. Immunophenotype of RBD-specific Bmem in individual donors by age.** (A) Absolute numbers and (B) Relative composition of RBD-specific Bmem in 30 BNT162b2 vaccinated individuals post dose 1. (C) Absolute numbers and (D) Relative composition of RBD-specific Bmem post dose 2 of the BNT162b2 vaccine. Dotted lines in A and C represent median RBD-specific Bmem numbers.

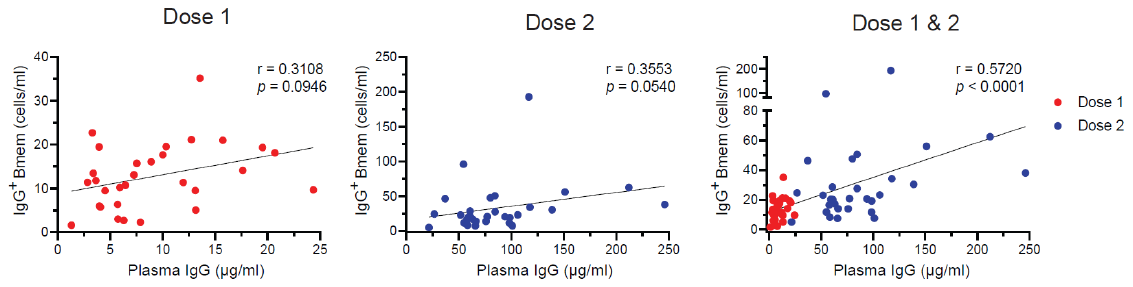

**Supplementary Figure 3. Correlation between plasma IgG and IgG^+^ RBD-specific Bmem.** Correlation between plasma IgG and IgG^+^ RBD-specific Bmem following dose 1, -dose 2 and dose 1 and 2. Trend lines depict linear correlations, statistics were performed using the nonparametric Spearman’s rank correlation.

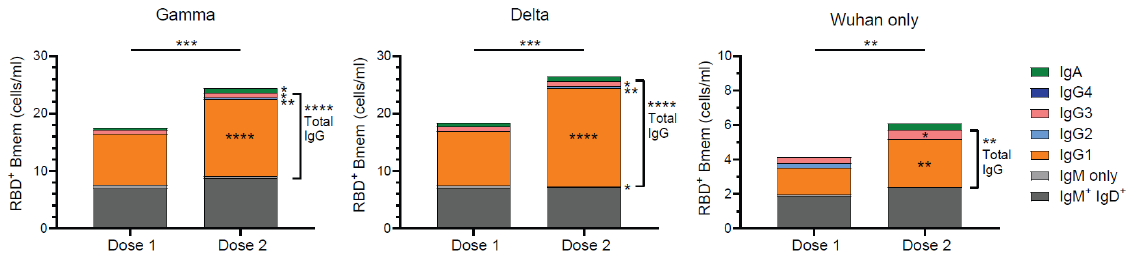

**Supplementary Figure 4. Immunophenotype of variant binding RBD-specific Bmem.** Absolute number of Ig switched total and RBD-specific Bmem that also bind the RBD of Delta, Gamma or Wuhan only. Wilcoxon matched-pairs signed rank test, * p < 0.05, ** p < 0.01, *** p < 0.001, **** p < 0.0001.
